# To Smooth or not to Smooth: Enhancing Specificity While Maintaining Sensitivity

**DOI:** 10.1101/2022.06.11.495739

**Authors:** Eileen Luders, Robert Dahnke, Christian Gaser, Alzheimer’s Disease Neuroimaging Initiative

**Affiliations:** School of Psychology, University of Auckland, Auckland, New Zealand; Department of Women’s and Children’s Health, Uppsala University, Uppsala, Sweden; Laboratory of Neuro Imaging, School of Medicine, University of Southern California, Los Angeles, USA; Department of Psychiatry and Psychotherapy, Jena University Hospital, Jena, Germany; Department of Neurology, Jena University Hospital, Jena, Germany; German Center for Mental Health (DZPG)

**Author notes:** Correspondence should be addressed to: Christian Gaser, Ph.D. Data used in preparation of this article were obtained from the Alzheimer’s Disease Neuroimaging Initiative (ADNI) database (adni.loni.usc.edu). As such, the investigators within the ADNI contributed to the design and implementation of ADNI and/or provided data but did not participate in analysis or writing of this report. A complete listing of ADNI investigators can be found at: http://adni.loni.usc.edu/wp-content/uploads/how_to_apply/ADNI_Acknowledgement_List.pdf.

**Keywords:** spatial smoothing, TFCE, neuroscience, non-parametric statistics, MRI, human brain mapping

## Abstract

Traditionally, when conducting voxel- or vertex-wise analyses in neuroimaging studies, it seemed imperative that brain data are convoluted with a Gaussian kernel, a procedure known as “spatial smoothing”. However, we suggest that – under certain conditions – smoothing may be omitted for the benefit of an improved regional specificity. We demonstrate the suitability of this omission by combining high-dimensional spatial registration and threshold-free cluster enhancement (TFCE) in a sample of 754 brains. Our findings revealed that, without smoothing, it is possible to capture brain atrophy within the hippocampal complex while dissociating neighboring areas (cornu ammonis, dentate gyrys, subiculum, and amygdala). In contrast, the traditional smoothing step would result in a single hippocampal cluster (the larger the smoothing kernel, the lower the specificity). Supplemental analyses not only varying the size of the smoothing kernel, but also the size of the sample, the signal-to-noise ratio, as well as the accuracy of the spatial registration confirm that no smoothing (or less smoothing) leads to increased specificity while maintaining sensitivity, at least for small-scale structures (e.g., hippocampus and amygdala). Nevertheless, classic analyses based on smoothed data will continue to provide important insights, especially for large-scale structures (e.g., cortical regions).

## Introduction

Spatial smoothing is a key preprocessing step in the field of brain mapping, where image data are convoluted with a (Gaussian) kernel before they are statistically analyzed. Smoothing ensures that data are normally distributed (a pre-requisite for conducting parametric tests and for the theory of Gaussian Random Fields), accounts for any remaining anatomical variations across brain scans after the spatial registration, and improves the signal-to-noise ratio (which increases the sensitivity of the statistical analysis). However, smoothing ultimately also poses a problem as it diminishes the regional specificity of the analysis outcomes (the larger the smoothing kernel, the lower the specificity). As a result, when generating the statistical map, significance clusters often spread across anatomical boundaries (the larger the smoothing kernel, the higher the spread). That is, valuable regional information originally inherent in the acquired brain image gets lost during the smoothing procedure and spatial accuracy decreases.

In other words, there are considerable drawbacks to smoothing and the question arises whether we can do without spatial smoothing under certain conditions, such as (I) when the statistical approach does not require a normal distribution of data; (II) when brain images are spatially aligned with high accuracy, (III) and when statistical degrees of freedom and/or anticipated effect sizes are sufficiently large. Ensuring that all three conditions are met may seem challenging, but is possible considering new developments in the field of human brain mapping. More specifically, with respect to condition I, there are numerous statistical approaches based on non-parametric inference (i.e., data do not need to be normally distributed), among them threshold-free cluster enhancement (TFCE; Smith and Nichols, 2009). TFCE has the additional benefit of being relatively sensitive (which is also relevant for condition III), a result of integrating cluster information (i.e., cluster size significance) with voxel information (i.e., peak voxel significance). With respect to condition II, there are a multitude of spatial registration algorithms which allow for an almost perfect overlap and spatial correspondence across brain images (Klein et al., 2009), among them high-dimensional warping (Ashburner and Friston, 2011; Avants et al., 2008; Klein et al., 2009). With respect to condition III, there is an ever increasing pool of large-scale databases, often containing hundreds or even thousands of brain scans, among them the Alzheimer’s Disease Neuroimaging Initiative (ADNI; https://adni.loni.usc.edu/) or the UK Biobank (https://www.ukbiobank.ac.uk/). Larger samples result in higher degrees of freedom, which increases the sensitivity and power of the statistical analysis. However, even within small samples, effect sizes might be sufficiently large (e.g., when assessing the impact of neurodegenerative diseases on the brain) to manifest as significant effects.

Here, we leveraged a large dataset (n=754) from ADNI to test whether analyzing unsmoothed data affords an adequate sensitivity and leads to an improved regional specificity compared to analyzing smoothed data using standard kernel sizes of 2 mm, 4 mm, and 6 mm full width at half maximum (FWHM). For this purpose, we conducted a voxel-based morphometry (VBM) analysis, in association with TFCE (Smith and Nichols, 2009) and high-dimensional warping (Ashburner and Friston, 2011), comparing voxel-wise gray matter between four subgroups: healthy controls (CTL; n=218), individuals with stable Mild Cognitive Impairment (sMCI; n=222), individuals with progressive MCI (pMCI; n=130), and individuals with Alzheimer’s disease (AD; n=184).

## Materials and Methods

### Study Sample

Data used in the preparation of this article were obtained from the Alzheimer’s Disease Neuroimaging Initiative (ADNI) database (adni.loni.usc.edu). More specifically, we used the T1-weighted MRI data from ADNI 1, where baseline MRI data and test scores in selected cognitive scales (i.e., Mini-Mental State Examination [MMSE]) were available. Altogether, the sample contained 754 individuals, who were classified into four groups as (I) healthy controls (CTL) if they were cognitively healthy at baseline as well as at three-year follow-up (n = 218; males/females = 111/107, mean/SD age = 76.03/5.04 years, mean/SD MMSE = 29.13/1.01); (II) individuals with stable MCI (sMCI), if they were diagnosed with MCI at baseline as well as at three-year follow-up (n = 222, males/females = 145/77, mean/SD age = 75.51/7.32 years, mean/SD MMSE = 27.20/1.79), (III) individuals with progressive MCI (pMCI) if they were diagnosed with MCI at baseline and with AD at some point during the three-year follow-up without reversion to MCI (n = 130, males/females = 79/51, mean/SD age = 74.66/7.12 years, mean/SD MMSE = 26.68/1.75), and (IV) individuals with AD (AD), if they were diagnosed with AD at baseline as well as at three-year follow-up (n = 184, males/females = 95/99, mean/SD age = 75.28/7.56 years, mean/SD MMSE = 23.25/2.04).

### Main Analysis – Non-Smoothing versus Smoothing

All brain images were processed using the CAT12 toolbox (r1940; http://www.neuro.uni-jena.de/cat), as implemented in SPM12 (https://www.fil.ion.ucl.ac.uk/spm/software/spm12/). The default CAT12 settings were applied for bias field correction (Ashburner and Friston, 2005) and tissue classification, which was based on adaptive maximum a posteriori estimations (Rajapakse et al., 1997) and also accounted for partial volume effects (Tohka et al., 2004). As per CAT12 default, for the spatial registration, we applied geodesic shooting (Ashburner and Friston, 2011) but used a more detailed template with a resolution of 1 mm (default is 1.5 mm) and also generated the spatially registered gray matter segments with a resolution of 1 mm (default is 1.5 mm). The resulting gray matter segments underwent visual and automated quality checks and were finally multiplied generating four identical sets of data: The first set remained unsmoothed; the other three sets were smoothed with a Gaussian kernel of 2, 4, and 6 mm FWHM. To mitigate any segmentation artifacts on the gray/white border, an absolute gray matter threshold of 0.1 was applied to the unsmoothed data; the same mask was also applied to the other three sets (to avoid any bias due to different masks). The four datasets constituted the input for the statistical analysis: For each of the four datasets, we ran an ANCOVA with four groups (CTL, sMCI, pMCI, and AD), while removing the variance associated with total intracranial volume. The latter was calculated by adding the volumes of the gray matter, white matter, and cerebrospinal fluid segments in native space. To test for increasing atrophy across the four groups (CTL>sMCI> pMCI>AD), the following contrast was applied: 1.5 0.5 -0.5 -1.5. The statistical analysis was conducted using the non-parametric TFCE-toolbox (which is freely available at http://www.neuro.uni-jena.de/tfce) running 25,000 permutations and applying a threshold of 0.0001 that was family-wise error (FWE) corrected.

### Supplemental Analysis I – Geodesic Shooting vs. Other Spatial Registrations

In addition to testing how the choice of the smoothing kernel impacts the outcomes, we tested how the spatial accuracy of the registration alters the results. Basis for this supplemental analysis were five different sets of gray matter segments, all of them unsmoothed (0 mm FWHM) but each of them spatially registered in a different way: The first set was identical to the one described above; it was derived by geodesic shooting (Ashburner and Friston, 2011), a template resolution of 1 mm, and generating the registered images with a resolution of 1 mm (Shooting 1 mm). The second set was also derived by geodesic shooting, but with a resolution of 1.5 mm for template and resulting image (Shooting 1.5 mm), which is the default setting in CAT12. The third and fourth sets were derived by Diffeomorphic Anatomical Registration Through Exponentiated Lie Algebra (DARTEL; Ashburner, 2007), using 1 mm and 1.5 mm, respectively for template and resulting image (Dartel 1 mm and Dartel 1.5 mm). The fifth set was derived by applying the standard procedure as implemented in SPM12 (SPM12) based on the unified segmentation (Ashburner and Friston, 2005). For each of the five datasets (in reality four additional datasets), we ran an ANCOVA with four groups (CTL, sMCI, pMCI, and AD), exactly as described above for the main analysis.

### Supplemental Analysis II – Specificity and Sensitivity of TFCE vs. Conventional T Statistics

Finally, we also systematically investigated how specificity and sensitivity achieved via TFCE (as well as conventional T statistics) vary in dependence of the size of the smoothing kernel, the size of the sample, as well as the noisiness of the data. For this purpose, we simulated data using an atlas of the left hippocampus and its subfields (Winterburn et al., 2013) as well as of the left amygdala (Treadway et al., 2015), as implemented in the CoBrA atlas, which is provided with CAT12 (Gaser et al., 2022). More specifically, we created a 3D dataset in which half of the images contained no signal, while the other half simulated a signal in a specific subregion (i.e., CA1, subiculum, CA4/dentate gyrus, CA2/CA3, or amygdala). This dataset constitutes the ground truth.

To investigate the impact of impact of signal-to-noise, two different levels of signal-to-noise ratios (SNR) were added to the aforementioned dataset (SNR = 1 and 5), in accordance with prior simulations related to TFCE (Smith and Nichols, 2009). To investigate the impact of spatial smoothing, the 3D dataset was convoluted with different kernel sizes (FWHM = 0, 2, 4 and 6 mm FWHM), in accordance with our main analysis. To investigate the impact of sample size, four 3D datasets were generated (n = 60, 120, 240 and 480).

The resulting 32 datasets were then used to detect significant effects using a two-sample T-test by comparing data with and without signal. Here, we ran both TFCE and conventional T statistics to be able to compare the outcomes between both approaches. In either case, 5,000 permutations were used to randomly shuffle the data to estimate a threshold of p<0.05 that was FWE-corrected. Finally, the statistical effects were compared to the ground truth (as based on the original 3D dataset) to estimate the specificity and sensitivity of TFCE and T statistics, respectively.

## Results

### Main Analysis

As shown in **Figure 1**, the most pronounced effects (red clusters) were detected bilaterally in the hippocampus, amygdala, as well as the parahippocampal and entorhinal gyrus, with significantly more gray matter in CTL than in sMCI, pMCI, and AD. While these effects were evident in both unsmoothed (FWHM = 0 mm) and smoothed data (FWHM = 2 mm, 4 mm, 6 mm), findings were spread wider and clusters bled over anatomical boundaries in smoothed data (the larger the FWHM, the larger the spread). In contrast, when using unsmoothed data, there was no bleeding over. In fact, the resulting statistical map seemingly preserved the regional information of the initial brain scans and it is possible to discriminate between significance clusters pertaining to known subfields of the hippocampal complex and adjacent regions, such as the cornu ammonis (CA), the dentate gyrus (DG), the subiculum (SUB), and the amygdala (AMY). **Supplemental Figure 1** provides group-specific density plots (HC, sMCI, pMCI, AD) for the peak voxels derived at those distinct clusters (CA, DG, SUB, AMY) using unsmoothed data (FWHM = 0 mm).

**Figure 1.**
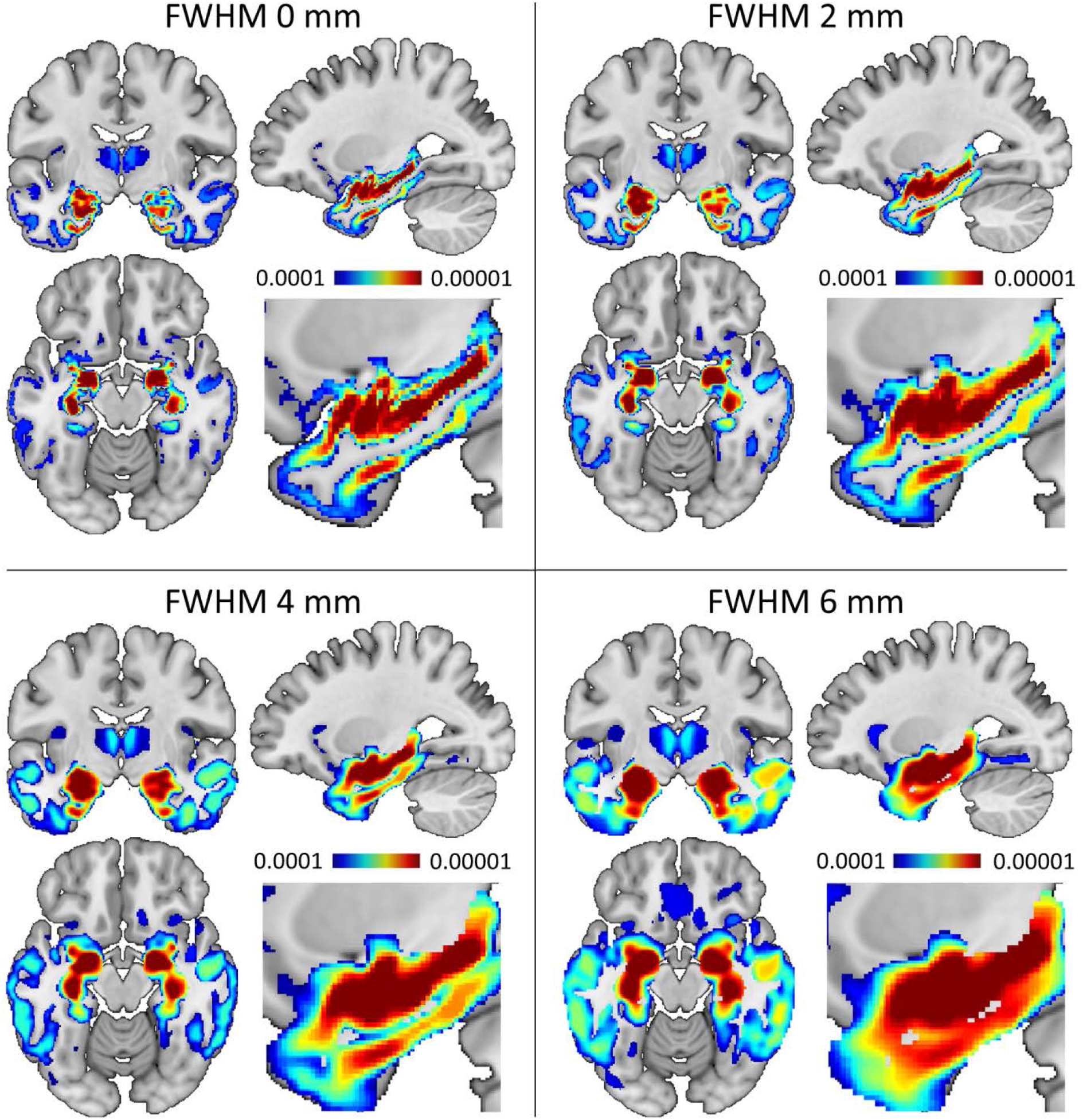
Significance maps with and without smoothing. All outcomes were derived for contrast CTL>sMCI>pMCI>AD at p<0.0001 using TFCE and FWE. Unsmoothed data (FWHM = 0 mm) and data smoothed with different kernel sizes (FWHM = 2 mm, 4 mm, and 6 mm. Orthogonal slices at x; y; z = -28 mm; -11 mm; -15 mm. The close-up sagittal slice (x = -28) provides a magnified view of the hippocampal complex. The color bar encodes significance (p) ranging between 0.0001 (blue) and 0.00001 (red). The highest regional specificity is achieved without smoothing (FWHM = 0 mm).

### Supplemental Analysis I

As shown in **Supplemental Figure 2**, the highest regional specificity and sensitivity is achieved by applying geodesic shooting and using a voxel size of 1 mm, both for the template and the spatially registered images (Shooting 1mm). The lowest regional specificity and sensitivity is achieved by applying SPM12’s standard spatial registration (SPM12). The other three approaches – geodesic shooting with a resolution of 1.5 mm (Shooting 1.5mm) as well as DARTEL with a resolution of 1 mm (Dartel 1mm) and 1.5 mm (Dartel 1.5mm) yielded similar maps.

### Supplemental Analysis II

Figure 2. illustrates the specificity for TFCE in dependence of varying kernel sizes (0 mm vs. 2 mm vs. 4 mm vs. 6 mm) in a sample of N=480 and a signal-to-noise ratio of 1. **Supplemental Figure 3** illustrates specificity and sensitivity for TFCE as well as for T statistics, in dependence of varying kernel sizes (0 mm vs. 2 mm vs. 4 mm vs. 6 mm), sample sizes (N=60 vs. N=120 vs. N=240 vs. N=480), and signal-to-noise ratios (1 vs. 5). As shown in **Figure 2**, the specificity of TFCE varies in dependence of the kernel size, with the highest specificity (values close to 0 on the x-axis) without smoothing (FWHM = 0 mm). Moreover, as demonstrated in **Supplemental Figure 3**, the sensitivity of TFCE is high (values close to 1 on the y-axis) regardless of the smoothing kernel. It is also relatively robust against variations in sample size and/or signal-to-noise. In contrast, the sensitivity of the T statistics is not only impacted by the size of the smoothing kernel but also by the size of the sample as well as the signal-to-noise ratio. However, just as it was observed for TFCE, the specificity of the T statistics increases with decreasing size of the smoothing kernel.

**Figure 2.**
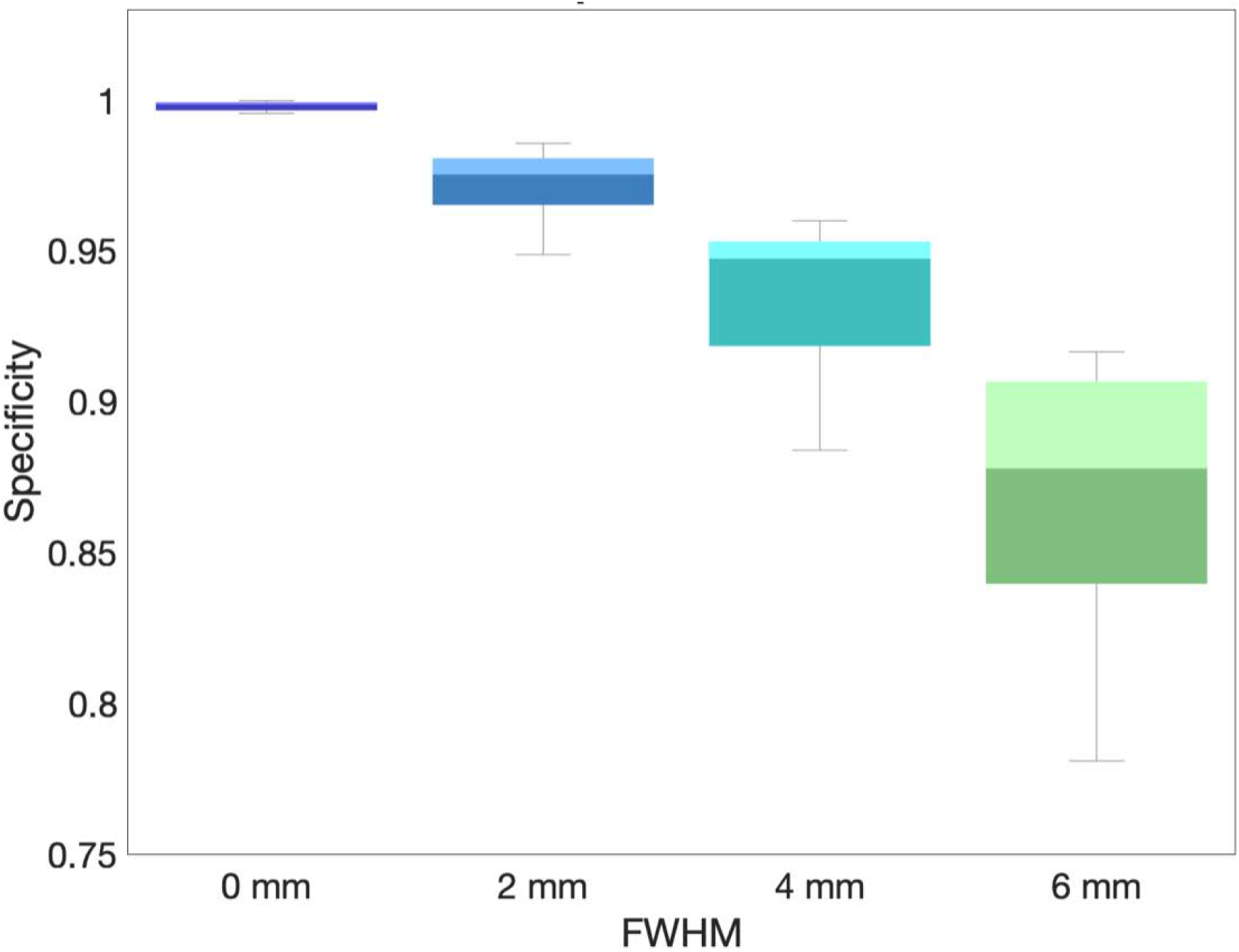
Specificity with and without smoothing. The x-axis shows the level of smoothness (FWHM = full width at half maximum), the y-axis the specificity. The whiskers indicate the 1.5 interquartile range; the change in color within each box the median. The highest specificity (close to 1.0) is achieved without smoothing (FWHM = 0 mm).

## Discussion

The main purpose of our study was to test the effect of omitting spatial smoothing, an essential step in traditional neuroimaging analyses. The focus of the study was on voxel-wise gray matter in MCI and AD, clinical conditions known to affect the hippocampus. The most pronounced effects (i.e., significantly less gray matter in MCI / AD patients compared to healthy controls) were detected within the hippocampal complex, which is well in line with other reports in the literature (Miller et al., 2012; Poulin et al., 2011; Turner and Geyer, 2014). While the aforementioned effects were evident in both smoothed and unsmoothed data, the statistical maps based on unsmoothed data provided a much higher regional specificity. That is, there was no bleeding over anatomical boundaries and it was even possible to discriminate between known subfields of the hippocampal complex and adjacent regions.

Thus, neuroimaging studies may benefit from omitting the smoothing step or from reducing the degree of smoothing, at least under certain conditions: (a) when brain images are spatially aligned with high accuracy (we used a high-dimensional spatial registration approach and demonstrated the advantage of geodesic shooting over DARTEL and SPM12); (b) when the statistical approach does not require a normal distribution of data (we used permutation testing and demonstrated the advantage of TFCE over conventional T statistics, especially at small sample sizes and lower signal-to-noise ratios); and (c) when anticipated effect sizes and/or degrees of freedom are large enough (we focused on a clinical condition known to impact the brain and we used a sample size of n=754). However, simulations suggest that, even at smaller sample sizes, unsmoothed data afford a sufficient sensitivity (i.e., effects become significant) when analyzed using TFCE statistics.

Skipping the spatial smoothing step may seem radical, but evidence for the suitability of this approach is increasing (Turner and Geyer, 2014). In fact, there seems to be a shift in attitude towards smoothing in the field of human brain mapping, complementing traditional views that spatial smoothing is absolute necessary with more differentiated opinions (Brodoehl et al., 2020; Coalson et al., 2018; Glasser et al., 2018; Glasser et al., 2016a; Glasser et al., 2016b; Turner and Geyer, 2014). The current findings are in close resemblance with this shift in attitude. However, while we demonstrate the suitability of this approach for subregions of the hippocampus and amygdala (i.e., small structures) and in clinical conditions with profound impact on brain anatomy (i.e., tissue loss), smoothing may still be warranted for larger structures and/or in studies where effects on the brain are minute. So, rather than proposing the omission of the smoothing step as a universal remedy, we encourage neuroscientists to conduct both analysis streams – with and without smoothing (or at a reduced degree of smoothing) – and see if the gain in regional specificty is offset by a loss in sensitivity or not. No losses or only mall losses in sensitivity may warrant omission of the smoothing step.

## Acknowledgments

Data collection and sharing for this project was funded by the Alzheimer’s Disease Neuroimaging Initiative (ADNI) (National Institutes of Health Grant U01 AG024904) and DOD ADNI (Department of Defense award number W81XWH-12-2-0012). ADNI is funded by the National Institute on Aging, the National Institute of Biomedical Imaging and Bioengineering, and through generous contributions from the following: AbbVie, Alzheimer’s Association; Alzheimer’s Drug Discovery Foundation; Araclon Biotech; BioClinica, Inc.; Biogen; Bristol-Myers Squibb Company; CereSpir, Inc.; Cogstate; Eisai Inc.; Elan Pharmaceuticals, Inc.; Eli Lilly and Company; EuroImmun; F. Hoffmann-La Roche Ltd and its affiliated company Genentech, Inc.; Fujirebio; GE Healthcare; IXICO Ltd.; Janssen Alzheimer Immunotherapy Research & Development, LLC.; Johnson & Johnson Pharmaceutical Research & Development LLC.; Lumosity; Lundbeck; Merck & Co., Inc.; Meso Scale Diagnostics, LLC.; NeuroRx Research; Neurotrack Technologies; Novartis Pharmaceuticals Corporation; Pfizer Inc.; Piramal Imaging; Servier; Takeda Pharmaceutical Company; and Transition Therapeutics. The Canadian Institutes of Health Research is providing funds to support ADNI clinical sites in Canada. Private sector contributions are facilitated by the Foundation for the National Institutes of Health (www.fnih.org). The grantee organization is the Northern California Institute for Research and Education, and the study is coordinated by the Alzheimer’s Therapeutic Research Institute at the University of Southern California. ADNI data are disseminated by the Laboratory for Neuro Imaging at the University of Southern California.

## Funding

Carl Zeiss Foundation

Alexander von Humboldt Foundation

Marie Skłodowska-Curie Innovative Training Network (859890-SmartAge-H2020-MSCA-ITN-2019)

## Author contributions

Conceptualization: CG; Methodology: RD, CG; Investigation: CG; Visualization: EL, RD, CG; Writing—original draft: EL; Writing—review & editing: EL, RD, CG

## Data and materials availability

The sources for all data and analyses are provided in the main text.

## Supplemental Material

**Supplemental Figure 1.**
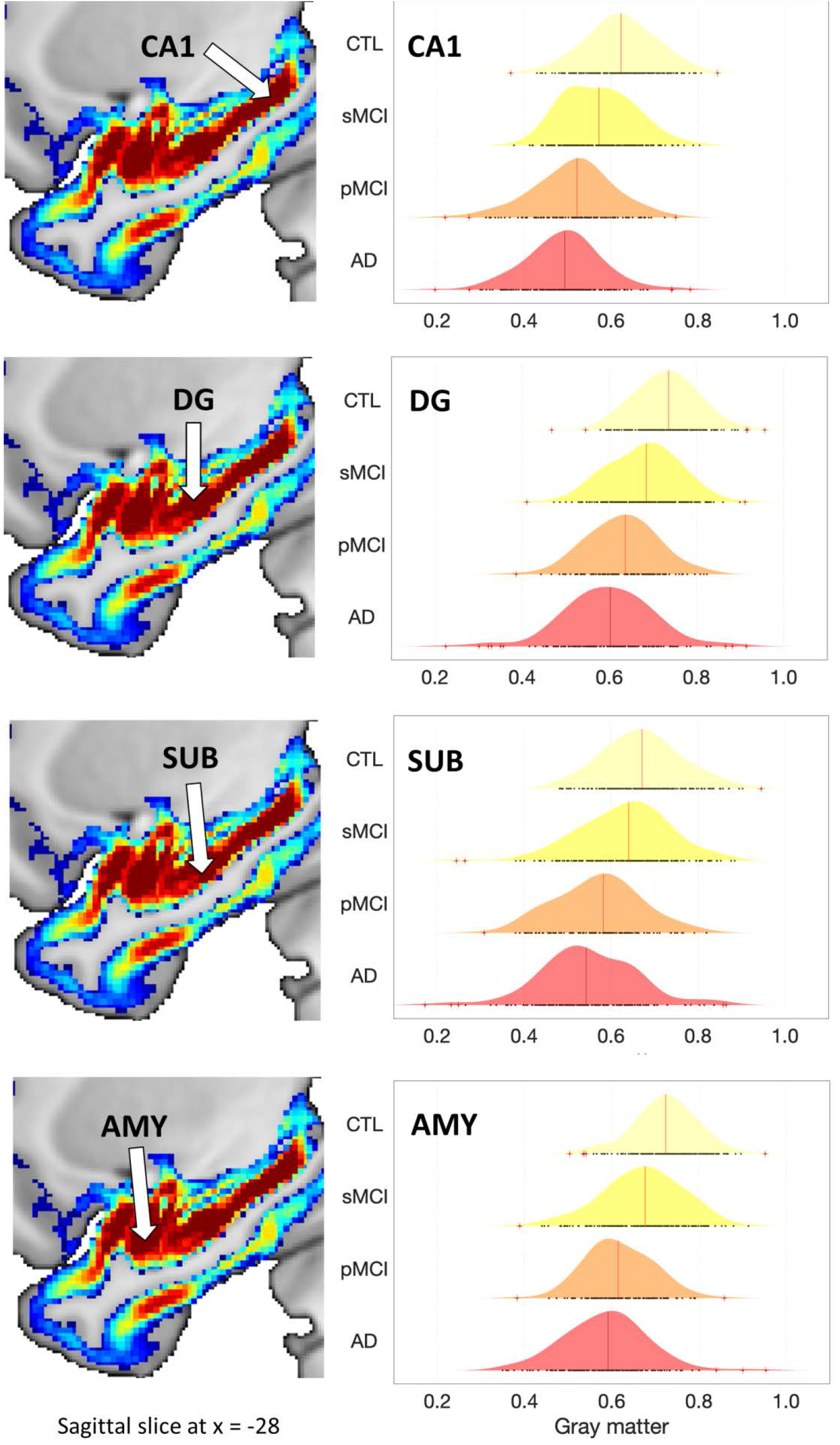
Density plots for regions of the hippocampal complex in unsmoothed data (FWHM = 0 mm). Right Panel: The x-axes show the amount of voxel-wise gray matter, the y-axes the four different groups: CTL = healthy controls, sMCI = stable mild cognitive impairment, pMCI = progressive mild cognitive impairment, and AD = Alzheimer’s disease. Left Panel: The arrows point to the approximate voxel for which the gray matter was plotted: CA1 = cornu ammonis (x; y; z = -28 -33 -11), DG = dentate gyrys (x; y; z = -28 -15 -20), SUB = subiculum (x; y; z = -28 -15 -25), and AMY = amygdala (x; y; z = -28 -6 -19). The significance map is identical with the one provided in Figure 1 (i.e., CTL>sMCI>pMCI>AD at p<0.0001 using TFCE and FWE).

**Supplemental Figure 2.**
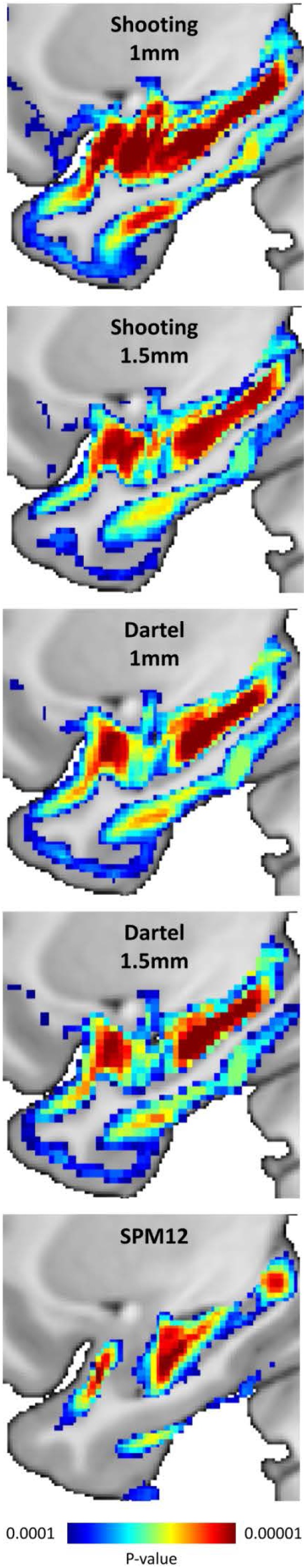
Impact of the Spatial Registration. The maps demonstrate the outcomes based on five different registration approaches with varying accuracy (highest to lowest accuracy, from top to bottom): geodesic shooting using a resolution of 1 mm (Shooting 1 mm); geodesic shooting using a resolution of 1.5 mm (Shooting 1.5 mm); Diffeomorphic Anatomical Registration Through Exponentiated Lie Algebra (DARTEL) using a resolution of 1 mm (Dartel 1mm); DARTEL using a resolution of 1.5 mm (Dartel 1.5mm); and (5) SPM’s standard procedure (SPM12). All maps were generated for contrast CTL>sMCI>pMCI>AD, based on unsmoothed data (FWHM = 0 mm) using TFCE and family-wise error (FWE) corrections at p<0.0001. The color bar encodes significance (p) ranging between 0.0001 (blue) and 0.00001 (red). Sagittal slice at x = -28.

**Supplemental Figure 3.**
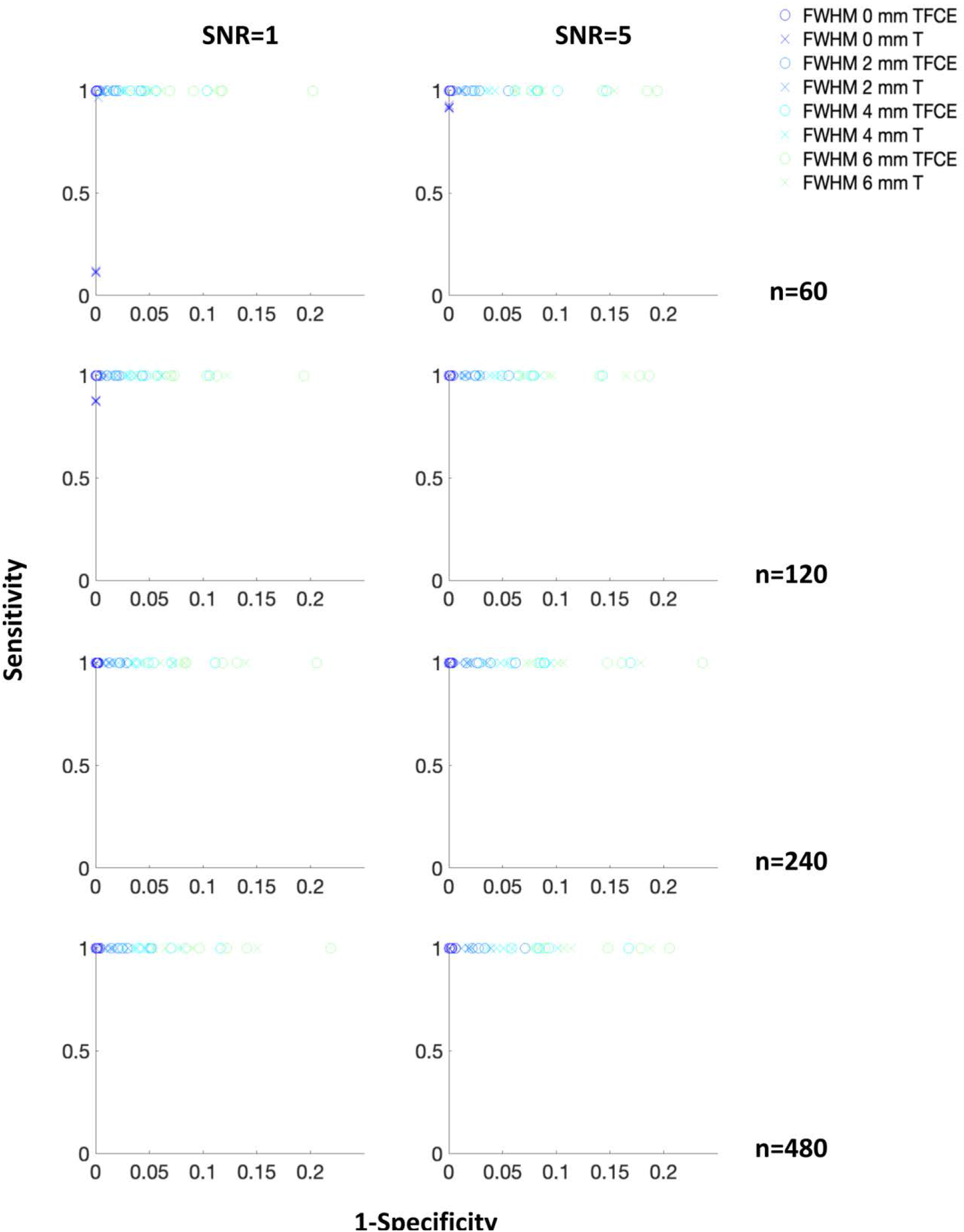
Impact of Kernel Size, Sample Size, and Signal-to-Noise. The x-axes show 1 minus the specificity (0 = highest specificity), the y-axes show the sensitivity (1 = highest sensitivity). The left and the right panels illustrate the outcomes for two different signal-to-noise ratios (SNR = 1 and 5); the rows illustrate the outcomes for four different sample sizes (n=60, 120, 240, and 480). The colored symbols (x = TFCE statistics; o = T statistics) indicate the outcomes in dependence of the applied full-width-at-half-maximum of the smoothing kernel (FWHM = 0, 2, 4 6 mm). TFCE = threshold-free cluster enhancement.

